# Genomic architecture of Shh dependent cochlear morphogenesis

**DOI:** 10.1101/665091

**Authors:** Victor Muthu, Alex. M. Rohacek, Yao Yao, Staci M. Rakowiecki, Alexander S. Brown, Ying-Tao Zhao, James Meyers, Kyoung-Jae Won, Shweta Ramdas, Christopher D. Brown, Kevin A. Peterson, Douglas J. Epstein

**Author notes:** Corresponding Author: Douglas J. Epstein, Ph.D., Professor and Vice Chair Department of Genetics Perelman School of Medicine University of Pennsylvania, Clinical Research Bldg., Room 463 415 Curie Blvd, Philadelphia, PA 19104.

## Abstract

The mammalian cochlea develops from a ventral outgrowth of the otic vesicle in response to Shh signaling. Mouse embryos lacking Shh or its essential signal transduction components display cochlear agenesis, however, a detailed understanding of the transcriptional network mediating this process is unclear. Here, we describe an integrated genomic approach to identify Shh dependent genes and associated regulatory sequences that promote cochlear duct morphogenesis. A comparative transcriptome analysis of otic vesicles from mouse mutants exhibiting loss (*Smo*^*ecko*^) and gain (*Shh-P1*) of Shh signaling revealed a set of Shh responsive genes partitioned into four expression categories in the ventral half of the otic vesicle. This target gene classification scheme provided novel insights into several unanticipated roles for Shh, including priming the cochlear epithelium for subsequent sensory development. We also mapped regions of open chromatin in the inner ear by ATAC-seq that, in combination with Gli2 ChIP-seq, identified inner ear enhancers in the vicinity of Shh responsive genes. These datasets are useful entry points for deciphering Shh dependent regulatory mechanisms involved in cochlear duct morphogenesis and establishment of its constituent cell types.

**SUMMARY STATEMENT:** An integrated genomic approach identifies Shh responsive genes and associated regulatory sequences with known and previously uncharacterized roles in cochlear morphogenesis, including genes that prime the cochlea for sensory development.

## INTRODUCTION

The mammalian cochlea derives from a ventral extension of the otic vesicle. Over the course of several days during embryonic development, this outgrowth undergoes a complex sequence of morphogenetic changes resulting in cochlear lengthening, coiling and differential patterning into sensory and nonsensory cell types that are essential for hearing (Wu and Kelley 2012; Basch et al., 2016; Montcouquiol and Kelley, 2019). Congenital malformations of the cochlea or defects in many of its constituent cell types are primary causes of hearing loss, emphasizing the importance of a thorough understanding of cochlear development (Jackler et al., 1987; Dror and Avraham, 2010; Schwander et al., 2010; Korver et al., 2017).

The organ of Corti is a specialized sensory structure for hearing in mammals that lines the length of the cochlear duct. It comprises a single row of inner hair cells (IHCs), three rows of outer hair cells (OHCs) and a variety of interspersed support cells that sit atop the basilar membrane. Sound waves are propagated through the cochlear duct by way of fluid motions that cause the basilar membrane to resonate at frequency dependent positions. OHCs enhance hearing sensitivity and frequency selectivity by amplifying basilar membrane vibrations in a feedback loop driven by OHC electromotility (Fettiplace 2017). Excitation of IHCs convert sound induced vibrations into electrochemical signals that are transmitted to the brain along auditory nerve fibers (Kazmierczak and Muller, 2012; Yu and Goodrich, 2014). Even slight deviations in the precise arrangement of sensory and non-sensory cell types in the organ of Corti can alter auditory perception (Montcouquiol and Kelley, 2019).

We previously described a critical function of the Sonic hedgehog (Shh) signaling pathway in promoting ventral identity within the otic vesicle that is necessary for the initiation of cochlear duct outgrowth (Riccomagno et al., 2002; Bok et al., 2007; Brown and Epstein, 2011). Mouse embryos lacking *Shh*, or carrying an <u>e</u>ar <u>c</u>onditional <u>k</u>nock<u>o</u>ut of *Smoothened* (*Foxg1cre; Smo*^*loxp*/−^, herein termed *Smo*^*ecko*^), an essential Shh signal transduction component, exhibit cochlear agenesis. We also classified several transcription factors (*Pax2, Otx2, Gata3*) with key roles in cochlear development as transcriptional targets of Shh signaling within the ventral otic epithelium (Brown and Epstein, 2011). Despite these advances, a detailed understanding of the mechanism by which Shh dependent transcription factors promote cochlear duct outgrowth remains unclear, in part, because the genes acting downstream in this transcriptional cascade have yet to be fully elucidated.

Shh regulates the expression of target genes through the Gli family of zinc finger containing transcription factors (Falkenstein and Vokes, 2014). In response to Shh signaling, transcription can be activated by the binding of full-length Gli proteins (Gli1, Gli2, Gli3) to cognate recognition sequences in the enhancers and promoters of target genes, often in conjunction with cooperating factors, or alternatively, by preventing the accumulation of a truncated Gli3 repressor (Bai et al., 2004; Peterson et al., 2012; Oosterveen et al., 2012; Oosterveen et al., 2013; Falkenstein and Vokes, 2014). Cochlear duct outgrowth is severely impaired in *Gli2*^−/−^;*Gli3*^−/−^ embryos and is not effectively restored in *Shh*^−/−^;*Gli3*^−/−^ double mutants (Bok et al., 2007). These genetic data suggest that Gli2 and Gli3 function primarily as transcriptional activators to promote the extension of the cochlear duct (Bok et al., 2007). However, as with the other Shh dependent transcription factors mentioned above, the inner ear specific targets of Gli2 and Gli3 remain unknown.

To identify novel targets of Shh signaling in the inner ear we performed RNA-seq on otic vesicles isolated from mouse mutants displaying a loss (*Smo*^*ecko*^) or gain (Shh-P1) in Shh signaling at E11.5, when cochlear outgrowth is first evident. We uncovered an intriguing set of Shh responsive genes with known and previously uncharacterized roles in cochlear morphogenesis. We also mapped regions of open chromatin in the inner ear by ATAC-seq (assay for transposase-accessible chromatin using sequencing) that, in combination with Gli2 ChIP-seq, identified inner ear enhancers in the vicinity of Shh responsive genes, several of which were functionally validated *in vivo* using a mouse transgenic reporter assay. This integrated genomic approach revealed several unexpected roles for Shh signaling, including transcriptional regulation of a set of genes that prime the medial wall of the cochlear duct for subsequent sensory development.

## RESULTS

### Screening for Shh responsive genes expressed during cochlear duct outgrowth

To identify a comprehensive set of genes regulated by Shh signaling at early stages of cochlear duct outgrowth we performed RNA-seq on otic vesicles isolated at E11.5 from two different mouse mutants and corresponding control littermates that were previously shown to exhibit loss (*Smo*^*ecko*^) and gain (*Shh-P1*) of Shh signaling in the otic epithelium (Riccomagno et al., 2002; Brown and Epstein, 2011). *Shh-P1* embryos display ectopic expression of *Shh* in the dorsal otic vesicle from a P1 transgene (Riccomagno et al., 2002). A total of 1,122 genes (581 downregulated, 541 upregulated) were differentially expressed between *Smo*^*ecko*^ and control otic vesicles (FDR≤0.05 and RPKM≥1.0), and 1,573 genes (670 downregulated, 903 upregulated) were differentially expressed between *Shh-P1* and control embryos (Fig. 1 A,B, Tables S1, S2). We intersected these datasets to uncover genes that were both downregulated in *Smo*^*ecko*^ and upregulated in *Shh-P1* (Shh activated genes), or upregulated in *Smo*^*ecko*^ and downregulated in *Shh-P1* (Shh repressed genes). This comparative transcriptome analysis revealed that Shh signaling is necessary and sufficient for the activation of 141 genes and repression of 77 genes in the otic vesicle at E11.5 (Fig. 1C-F, Fig. S1, Tables S3, S4).

**Figure 1.**
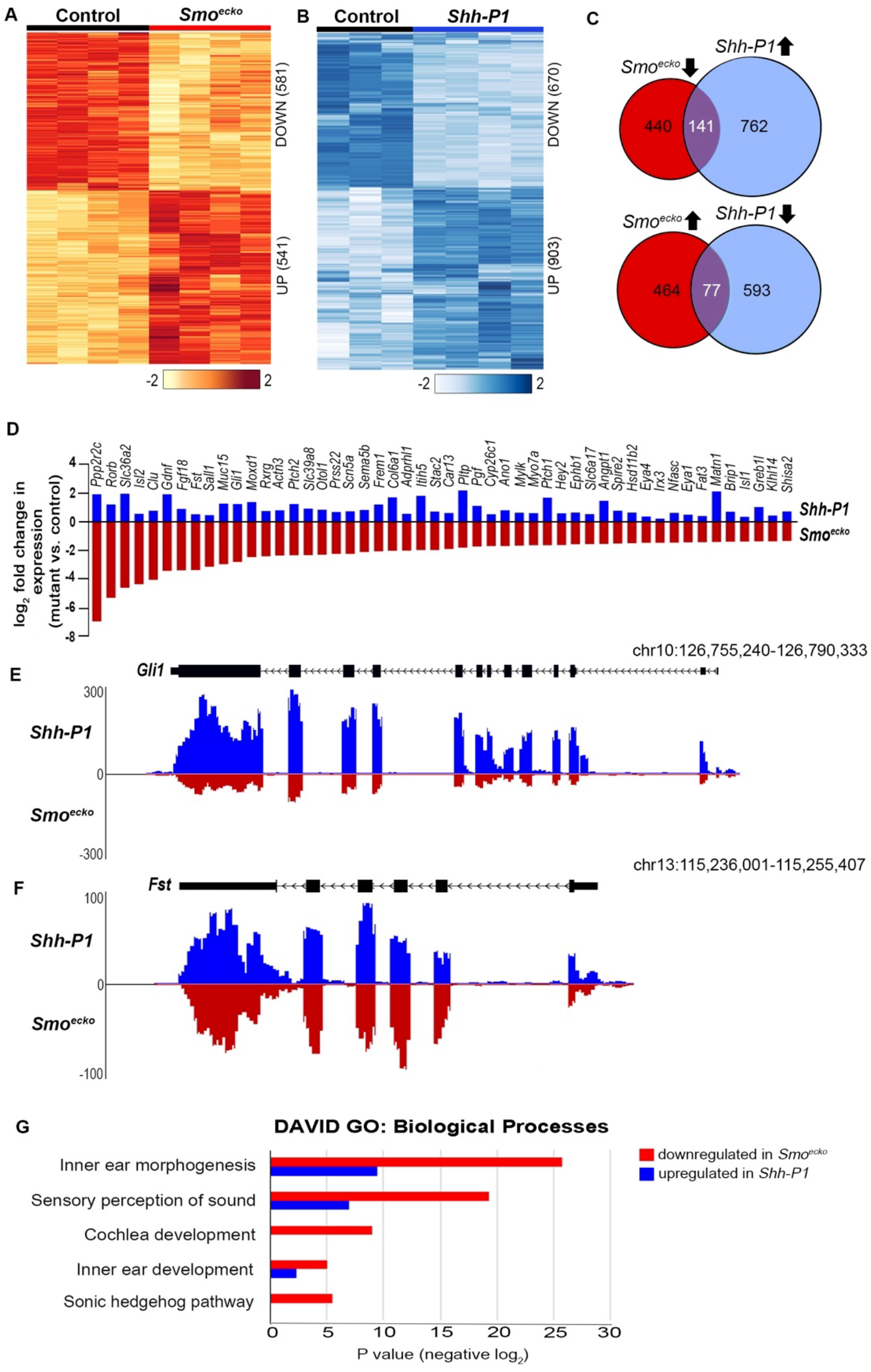
Differential expression profiling identifies Shh responsive genes in the inner ear. (A, B) Heat maps of differential RNA-seq profiles (log_2_ fold change) between (A) control and *Smo*^*ecko*^ (n=4), and (B) control (n=3) and *Shh-P1* (n=4) inner ears at E11.5 (FDR≥0.05 and RPKM≥1.0). (C) Intersection of differentially expressed genes in *Smo*^*ecko*^ (red) and *Shh-P1* (blue) inner ears identifies Shh activated (top) and Shh repressed (bottom) gene sets. (D) Top 50 Shh activated genes that show significant downregulation in *Smo*_*ecko*_ (red) and upregulation in *Shh-P1* (blue) inner ears (log_2_ fold change). (E, F) Normalized RNA-seq read counts in *Smo*^*ecko*^ (red) and *Shh-P1* (blue) mutants of two representative Shh activated genes, *Gli1* and *Fst*. (G) DAVID Gene Ontology term enrichment (Biological Processes) for gene sets that are downregulated in *Smo*^*ecko*^ (red) and upregulated in *Shh-P1* (blue) mutants.

Shh activated genes are significantly enriched in gene ontology (GO) terms associated with inner ear morphogenesis, sensory perception of sound, cochlear development and Hedgehog signaling activity (Fig. 1G). Of the top 50 Shh activated genes, 20 have documented inner ear expression in peer reviewed publications, including seven with established roles in cochlear development and/or auditory function (Fig. 1D). Genes in this category are likely to serve as Shh dependent regulators of cochlear development.

Shh repressed genes are also enriched for GO terms associated with inner ear morphogenesis (Fig. S1 and Table S4). Several of these genes (*Hmx2, Hmx3, Bmp4, Msx1, Msx2, Meis1, Meis2*) are expressed in dorsal regions of the otic vesicle, and/or have known roles in vestibular development (Wang et al., 2004; Chang et al., 2008; Sanchez-Guardado et al., 2011). These results suggest that Shh signaling within the otic epithelium may be required to prevent a subset of dorsal otic genes from being ectopically expressed in ventral regions of the otic vesicle.

Not all previously described Shh responsive genes in the inner ear were differentially expressed in both *Smo*^*ecko*^ and *Shh-P1* embryos. For instance, expression of the homeodomain transcription factor, *Otx2*, was downregulated five-fold in *Smo*^*ecko*^ mutants, but was unaltered in *Shh-P1* embryos (Tables S1, S2). On the other hand, some genes (e.g. *Six1*, *Eya1* and *Jag1*) that showed no change in mRNA transcript abundance between *Smo*^*ecko*^ and control littermates were significantly upregulated in *Shh-P1* embryos (Tables S1, S2). Hence, we considered any genes exhibiting loss and/or gain of expression in either *Smo*^*ecko*^ or *Shh-P1* embryos as candidate *Shh* responsive genes in the otic vesicle.

To validate the expression of candidate Shh responsive genes in the developing cochlear duct we selected 24 genes that were downregulated in *Smo*^*ecko*^ and/or upregulated in *Shh-P1* embryos for further analysis by in situ hybridization. Many of these genes (n=17) have known or predicted roles in cochlear development, including seven (*Emx2*, *Eya1*, *Eya4*, *Mpzl2*, *Pls1*, *Six1*, *Gata3*) that when mutated cause hearing loss in humans and/or mice (https://hereditaryhearingloss.org/recessive-genes; http://www.informatics.jax.org). The expression of seven additional genes (*Dsp, Pcdh11x, Brip1, Gas2, Fam107a, Slc39a8, Capn6*) had not previously been described in the otic vesicle.

Four genes (*Gli1*, *Pax2*, *Gata3* and *Otx2*), already characterized as Shh dependent transcription factors (Brown and Epstein, 2011), exhibit distinct patterns of expression in the otic vesicle at E11.5 that include broad ventral (*Gli1*), medial wall (*Pax2*), ventral tip (*Gata3*), and lateral wall (*Otx2*) domains (Fig. 2). Remarkably, all other genes selected for follow up analysis were expressed in one of these four Shh responsive regions. For instance, known (*Emx2*, *Eya1* and *Eya4*) and previously uncharacterized (*Dsp, Mpzl2 and Pcdh11x*) genes were broadly expressed in the ventral half of the otic vesicle in a similar pattern with *Gli1* (Fig. 2). Six genes (*Brip1, Car13, Gas2, Fam107a, Pls1 and Six1*) displayed overlapping expression with *Pax2* on the medial side of the otic vesicle (Fig. 2). Six other genes (*Ano1, Fst, Hey1, Jag1, Lin28b* and *Slc39a8*) were expressed in the ventral tip of the otic vesicle in a similar pattern to *Gata3*, including genes (*Hey1, Jag1, Lin28b*) implicated in prosensory development (Benito-Gonzalez and Doetzlhofer, 2014; Kiernan and Gridley, 2006; Golden et al., 2015). Finally, two genes (*Capn6, Rspo2*) were expressed on the lateral wall of the otic vesicle, comparable to *Otx2*. These results demonstrate the utility of our differential RNA-seq analysis for discovery of Shh responsive genes expressed in discrete ventral territories of the otic vesicle during the initial stages of cochlear duct outgrowth.

**Figure 2.**
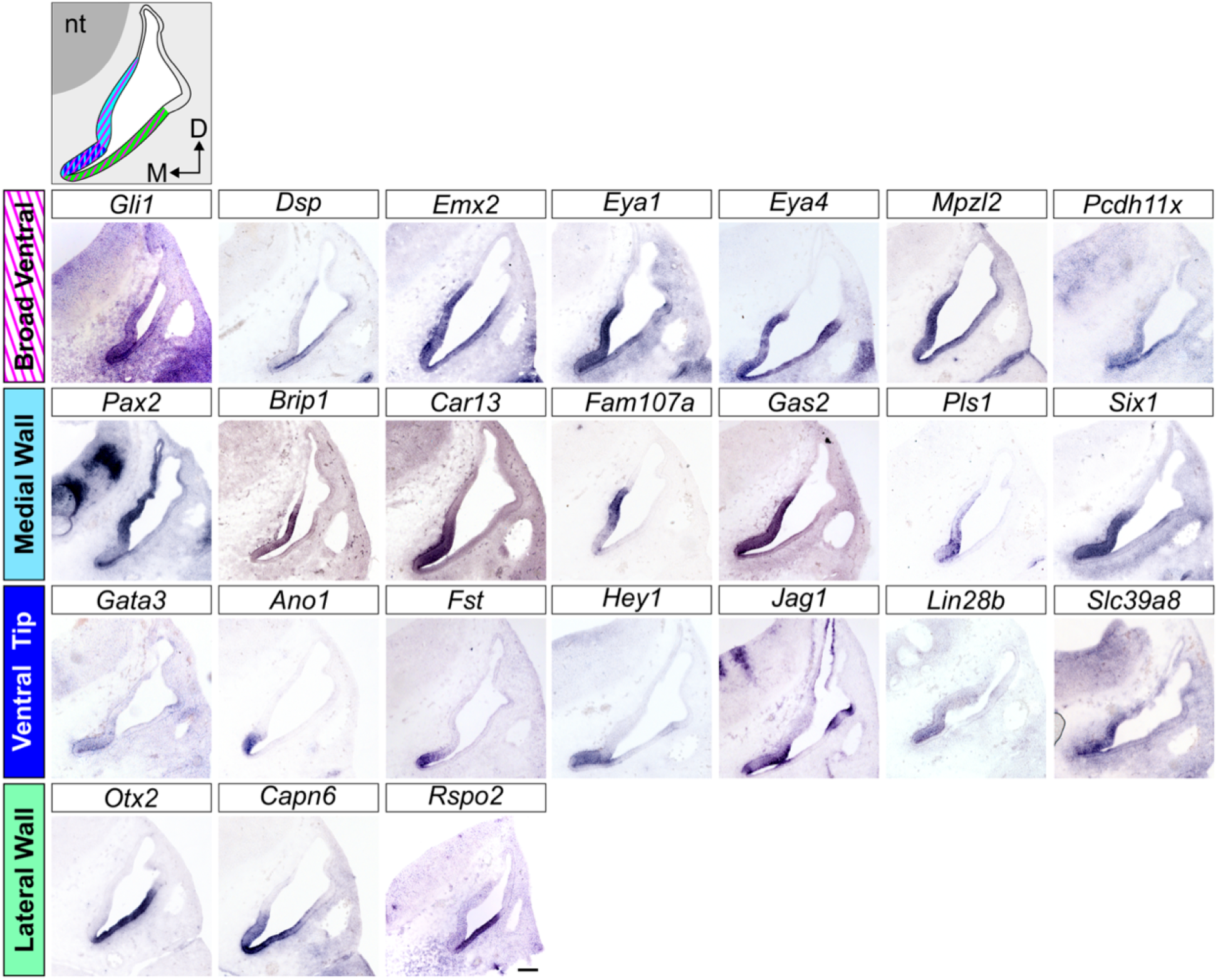
Classification of Shh responsive genes in wild type otic vesicles at E11.5. Schematic of a transverse section through the inner ear color coded to represent the four patterns of Shh responsive genes in broad ventral (magenta diagonal lines), medial wall (light blue), ventral tip (dark blue), and lateral wall (green) regions of the otic vesicle. Expression of Shh responsive genes as determined by in situ hybridization on wild type sections through the inner ear at E11.5 (n=3 replicates). Scale bar: 100 μm. Abbreviations: nt, neural tube; D, Dorsal; M, Medial.

Given that the otic vesicle fails to extend ventrally in *Smo*^*ecko*^ embryos at E11.5, the reduced expression of Shh responsive genes may be due to the loss of Shh signaling activity, or to the loss of ventral otic tissue. To discern between these two possibilities, we evaluated expression one day earlier, at E10.5, prior to the emergence of differences in otic vesicle morphology between *Smo*^*ecko*^ and control embryos (Fig. 3). Our earlier work revealed that expression of *Gli1*, *Pax2*, *Gata3*, and *Otx2* was absent from *Smo*^*ecko*^ otic vesicles at E10.5 (Brown and Epstein, 2011). Similarly, the majority of Shh responsive genes analyzed here showed abrogated otic expression in *Smo*^*ecko*^ mutants compared to control littermates at E10.5 (Fig. 3). It should be noted that compared to E11.5, the expression of several of these genes was weaker (*Dsp, Emx2, Mpzl2, Car13, Pls1, Hey1*) or not detected (*Fst, Pcdh11x*) in control embryos at E10.5.

**Figure 3.**
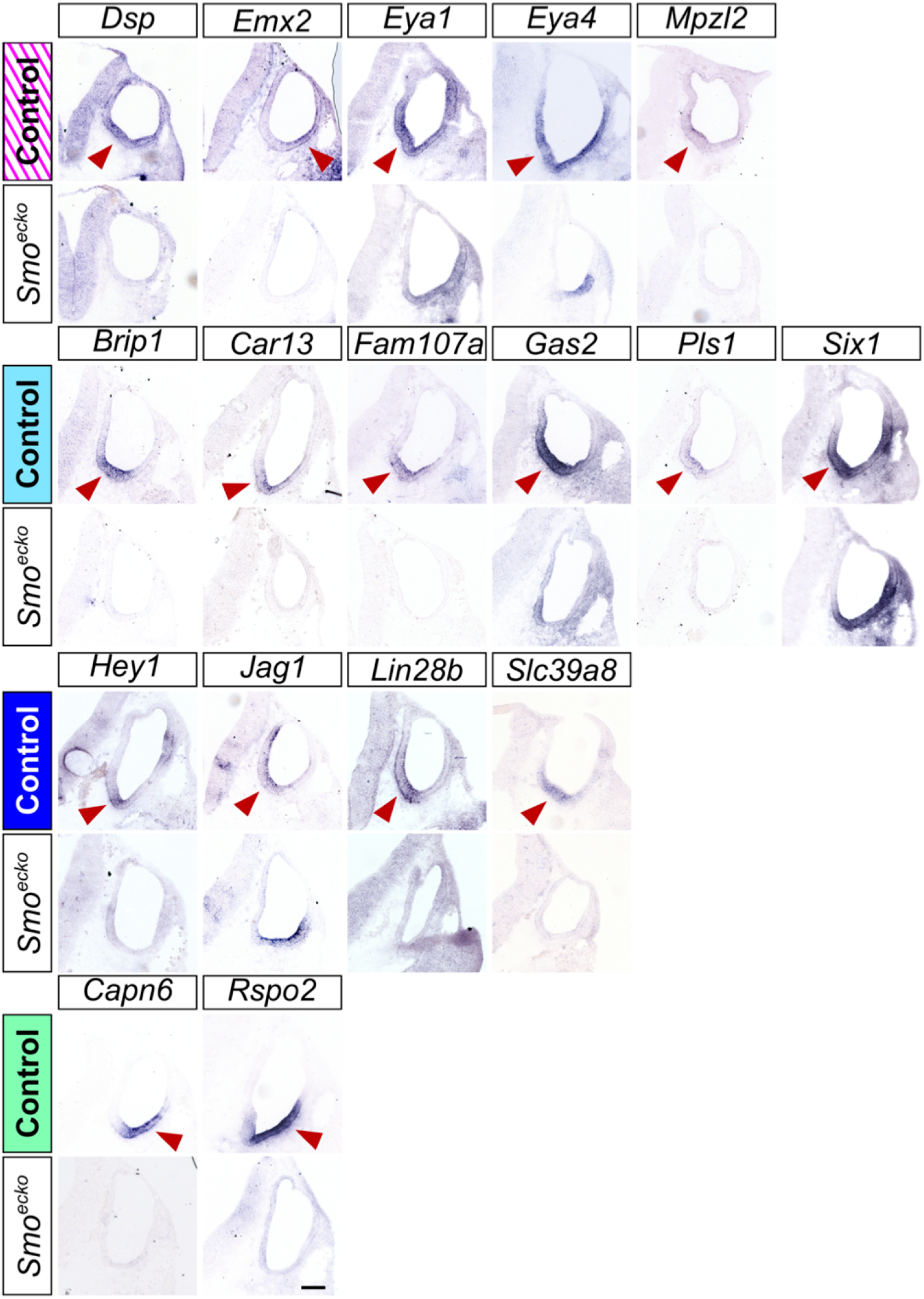
Shh responsive genes are misregulated in *Smo*^*ecko*^ embryos at E10.5. In situ hybridization of Shh responsive genes on transverse sections through control and *Smo*^*ecko*^ otic vesicles (n≥3 for all panels) at E10.5. Expression in control embryos (red arrowhead) for most genes is downregulated in *Smo*^*ecko*^ mutants with the exception of select genes (*Eya1, Eya4, Six1, Jag1*), which show maintained or ectopic expression on the lateral side of the otic vesicle. Scale bar: 100 μm.

The expression of *Six1* and *Jag1*, with defined roles in prosensory development, displayed more complex alterations in *Smo*^*ecko*^ mutants. These genes are prominently expressed on the ventromedial side of the otic vesicle in control embryos (Fig. 3), encompassing prospective sensory progenitors that begin to differentiate after E12.5 (Ruben, 1967; Chen and Segil, 1999). Interestingly, the expression of *Six1* and *Jag1* is flipped in *Smo*^*ecko*^ mutants, with loss of medial and gain of lateral otic staining (Fig.3). Two other genes (*Eya1* and *Eya4*) with broad ventral expression in the otic vesicle at E11.5 showed loss of medial and maintenance of lateral otic expression in *Smo*^*ecko*^ mutants (Fig. 3). Since Eya1 and Six1 form a transcriptional complex with Sox2 to regulate hair cell development (Ahmed et al., 2012), we evaluated Sox2 expression in *Smo*^*ecko*^ mutants, which also exhibited a medial to lateral switch in otic vesicle expression (Fig. S2). Not all genes expressed along the ventromedial wall showed flipped expression in *Smo*^*ecko*^ mutants, suggesting that this phenomenon may be specific for genes with prosensory function. Since Otx2 is required to repress sensory development on the lateral (nonsensory) side of the cochlear duct (Vendrell et al., 2015), we posit that the downregulation of *Otx2* in *Smo*^*ecko*^ embryos accounts for the derepression of genes with prosensory function on the lateral wall of the otic vesicle.

We also addressed the sufficiency of Shh to activate candidate target genes in the otic vesicle by evaluating their expression in *Shh-P1* embryos. The majority of genes (21/24) were ectopically expressed in broad dorsal (*Emx2, Eya1, Eya4, Mpzl2, Pcdh11x, Pax2, Pls1*) or lateral (*Dsp, Brip1, Car13, Gas2, Fam107a, Six1, Gata3, Ano1, Fst, Hey1, Jag1, Lin28b, S/c39a8*) regions of the otic vesicle in *Shh-P1* embryos at E11.5 (Fig. 4). Three genes (*Otx2, Capn6, Rspo2*) expressed in a ventrolateral otic domain did not show ectopic expression in *Shh-P1* embryos, suggesting that additional factors are required for their activation in conjunction with Shh. Taken together, these data identify a set of Shh responsive genes with known and potentially novel roles in early aspects of cochlear development.

**Figure 4.**
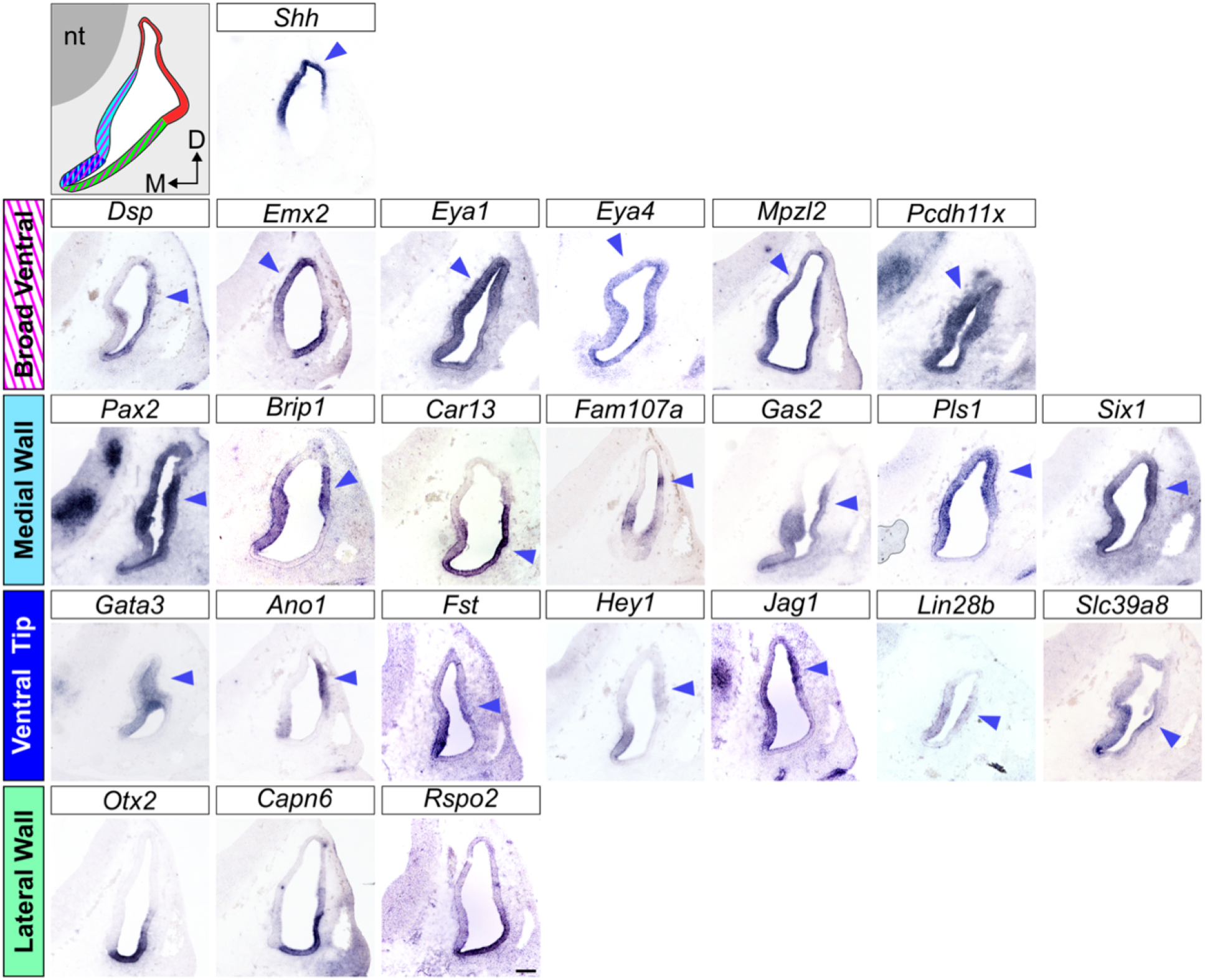
Shh responsive genes are ectopically expressed in *Shh-P1* embryos. Schematic of a transverse section through the inner ear of a *Shh-P1* embryo (E11.5) showing ectopic *Shh* expression in the dorsal otocyst (red). ln situ hybridization of *Shh* and Shh responsive genes on transverse sections through inner ears of *Shh-P1* embryos at E11.5 (n≥3 for all panels). Ectopic expression is indicated (blue arrowhead). Note that lateral wall genes (*Otx2*, *Capn6* and *Rspo2*) are not influenced by ectopic Shh signaling in *Shh-P1* embryos. Scale bar: 100 μm.

### ATAC-seq identifies active regulatory sequences in the otic vesicle

The expression of Shh responsive genes in the inner ear may be directly regulated by Gli2 and Gli3 (Bok et al., 2007), indirectly regulated by other Shh dependent transcription factors (e.g. *Pax2, Gata3, Otx2*), or by a combination of Shh dependent and independent transcription factors, as described in the spinal cord (Peterson et al., 2012; Oosterveen et al., 2012). As an entry point to deciphering the transcriptional mechanisms regulating Shh target gene expression in the inner ear, we performed ATAC-seq on chromatin isolated from wild type otic vesicles at E11.5. ATAC-seq profiles from four highly correlated biological replicates (peaks present in at least two samples) were merged, yielding 30,720 regions of open chromatin accessibility that map to intergenic (22.7%), intronic (24.3%), exonic (5.3%) and promoter (47.6%) regions of the genome (Fig. 5A).

**Figure 5.**
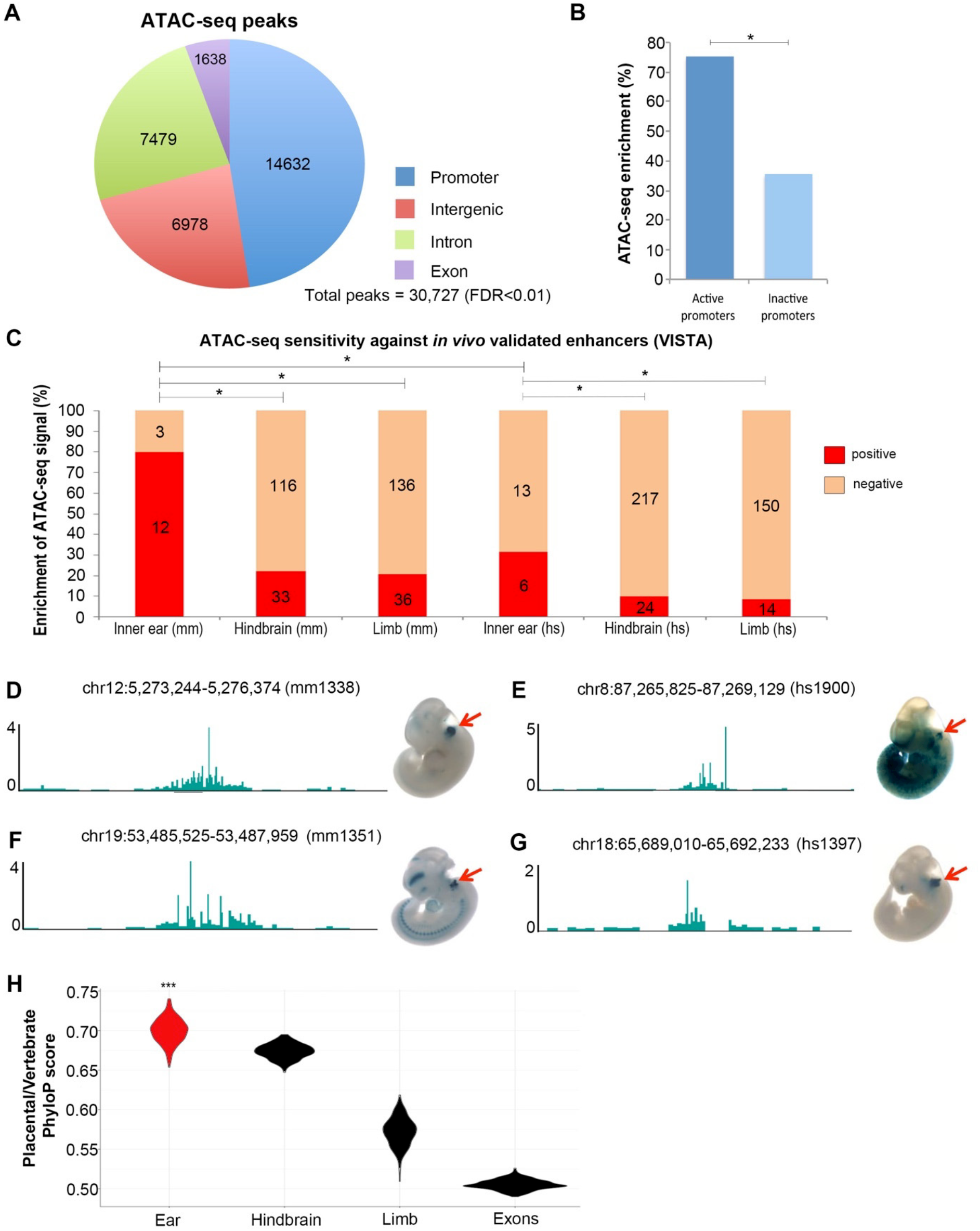
ATAC-seq identifies sites of open chromatin at inner ear regulatory sequences. (A) Genomic distribution of ATAC-seq peaks identified in the inner ear at E11.5 (FDR<0.01). (B) ATAC-seq signal enrichment on active versus inactive promoters at E11.5 (*p <0.05, Fisher’s exact test). (C) ATAC-seq signal enrichment on mouse (mm) and human (hs) enhancers active in the inner ear, hindbrain and limb from the VlSTA enhancer browser (*p<0.05, Fisher’s exact test). (D-G) ATAC-seq signal at representative mouse (D,F) and human (E,G) inner ear enhancers from the VlSTA enhancer browser. X-gal staining is detected in the otic vesicle (red arrow) of embryos (E11.5) carrying indicated reporter constructs. (H) Comparison of ATAC-seq sequence conservation (PhyloP score) from inner ear, hindbrain and limb between placental mammals and vertebrates. Exonic sequence from the inner ear was used as a deep conservation control (***p <2.2e-16, Kolmogorov-Smirnoff test). Error bars represent standard error of the mean.

Gene regulatory sequences typically reside within regions of open chromatin (Buenrostro et al., 2013; Vierstra and Stamatoyannopoulos, 2016). In agreement with this observation, promoters of actively transcribed genes in the inner ear (RNA-seq, RPKM≥1) were more likely to display ATAC-seq signal compared to promoters of inactive genes (RNAseq, RPKM<1) (Fig. 5B). ATAC-seq peaks were also selectively enriched on functionally validated mouse inner ear enhancers (80%) from the VlSTA Enhancer Database (Visel et al 2007), compared to hindbrain (22%) and limb (21%) enhancers (Fig. 5C,D,F). Similar ATAC-seq signal enrichment was observed for orthologous mouse sequence of human inner ear enhancers (32%) compared to hindbrain (10%) and limb (8.5%) enhancers, although levels are overall lower than for mouse enhancers (Fig. 5C,E,G).

Non-coding ATAC-seq peaks were also enriched in the vicinity of genes annotated for terms associated with inner ear gene expression and mouse phenotypes according to the GREAT analysis tool (Fig. S3; Mclean et al., 2010). Moreover, non-coding ATAC-seq peaks from the inner ear were specifically more conserved within placental mammals, as opposed to across more distantly related vertebrates, compared to other tissues, suggesting that evolutionary changes in gene regulatory sequences may underlie mammalian specific adaptations in inner ear morphology (Fig. 5H). Taken together, these findings suggest that the inner ear ATAC-seq dataset represents a robust resource for identifying functional regulatory sequences controlling gene expression in the developing otic vesicle.

### Discovery of Shh dependent inner ear enhancers through genomic integration

To identify Shh dependent regulatory sequences in the inner ear, we overlaid ATAC-seq and Gli2 ChIP-seq datasets (Fig. 6A). Unfortunately, it was not feasible to perform Gli2 ChIP-seq on isolated otic vesicles due to technical limitations with small scale tissue samples. Nevertheless, we took advantage of a Gli2 ChIP-seq dataset using embryonic mouse heads at E10.5, which included the otic vesicles (see methods). We found that 4% (605/14457) of intergenic and intronic ATAC-seq peaks overlapped with Gli2 occupied sites (Fig. 6B). Notably, gene set enrichment analysis demonstrated that Shh activated genes are significantly enriched around regions of open chromatin co-bound by Gli2 (FDR≥0.05), consistent with the premise that expression of these genes is directly regulated by Shh/Gli2 (Fig. 6C). Approximately, 20% (137/605) of overlapping ATAC-seq/Gli2 ChIP-seq sites also intersected with H3K27ac ChIP-seq peaks from E11.5 hindbrain (ENCODE project, ENCSR129LAP), a histone modification commonly associated with active enhancers (Fig. 6B). However, Shh activated genes were not enriched around overlapping genomic sites for these three signals, suggesting that the H3K27ac ChIP-seq dataset from hindbrain may not be a good predictor of Shh/Gli2 dependent enhancers in the inner ear (Fig. S4).

**Figure 6.**
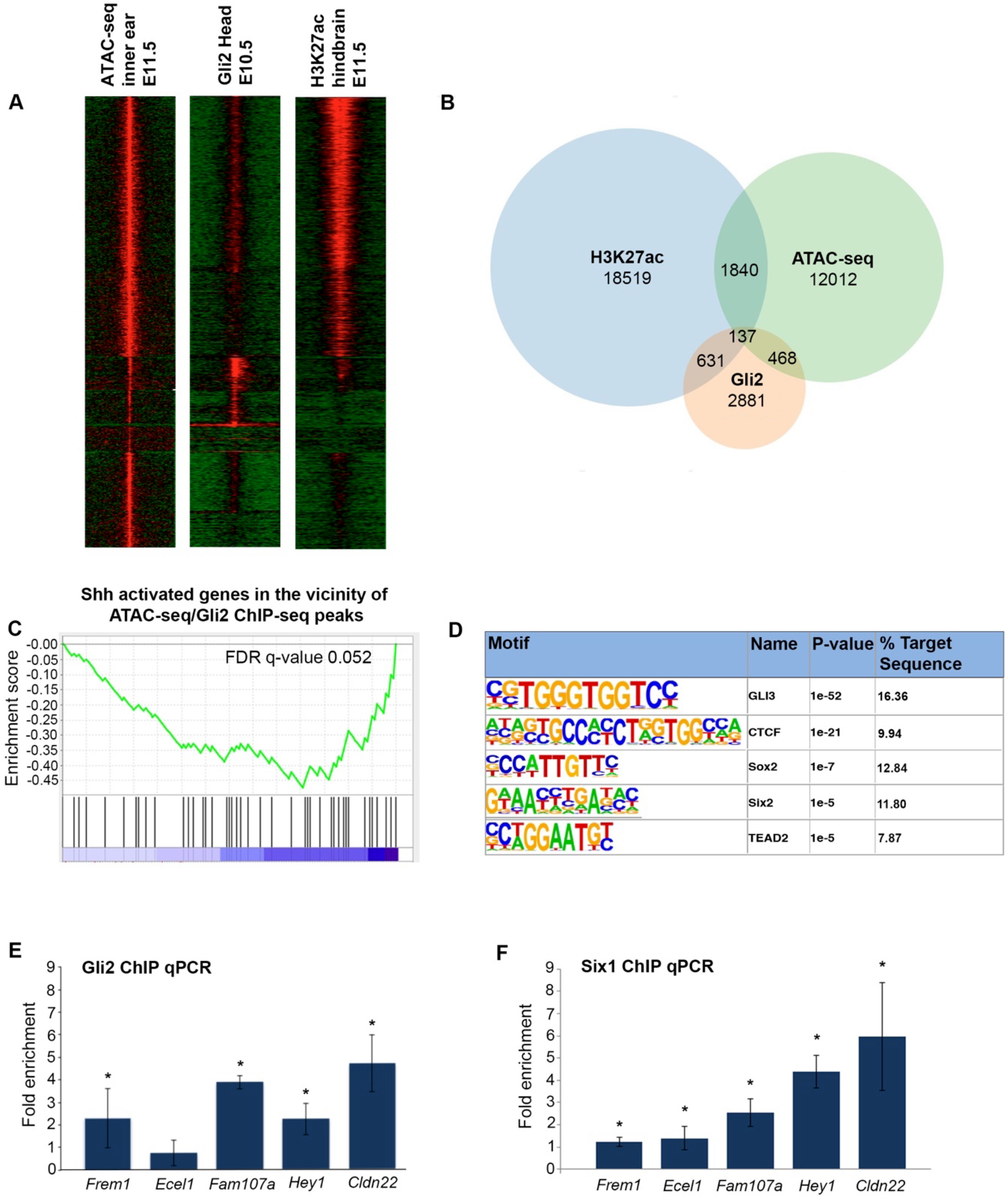
Identification of Shh dependent inner ear enhancers. (A) Heatmaps represent the overlap of ATAC-seq (E11.5 inner ear), Gli2 ChIP-seq (E10.5 head), and H3K27ac ChIP-seq (E11.5 hindbrain, ENCODE project, ENCSR129LAP) signals. (B) Venn diagram represents the intersection of ATAC-seq (intergenic and intronic), Gli2-ChIP-seq and H3K27ac-ChIP-seq sites. (C) Gene set enrichment analysis showing significant enrichment of Shh activated genes in the vicinity of overlapping ATAC-seq and Gli2 ChIP-seq peaks (FDR q-value=0.052). (D) HOMER motif enrichment analysis for intersected ATAC-seq and Gli2 ChIP-seq peaks. (E and F) ChIP-qPCR analyses showing significant co-recruitment of Gli2 and Six1 at candidate inner ear enhancers in the vicinity of Shh responsive genes (*p<0.05, Student’s t-test). Error bars represent standard error of the mean.

We next analyzed the DNA sequence at overlapping ATAC-seq/Gli2 ChIP-seq genomic regions for enrichment of motifs matching transcription factor binding sites (TFBS). As expected, the most over-represented motif matched the consensus binding sequence for Gli proteins (Fig. 6D). Other significantly enriched motifs included binding sites for CTCF, Sox2, Six and Tead family members. The presence of Sox2 and Six binding sites is particularly intriguing given that the activity of enhancers controlling expression of Shh/Gli target genes in the neural tube is often dependent on Sox2 (Peterson et al., 2012; Oosterveen et al., 2012), and both Sox2 and Six1 are essential for inner ear development (Zheng et al., 2003; Ozaki et al., 2004; Kiernan et al., 2005; Stevens et al., 2019).

*Six1*^−/−^ and *Smo*^*ecko*^ embryos display similar defects in cochlear duct outgrowth and misexpress several of the same genes in the otic vesicle (Ozaki et al., 2004; Brown and Epstein, 2011). Some of the phenotypic overlap between *Six1*^−/−^ and *Smo*^*ecko*^ may be attributed to altered *Six1* expression in *Smo*^*ecko*^ embryos (Fig. 3). A reciprocal downregulation in Shh signaling was not observed in *Six1*^−/−^ mutant ears (Ozaki et al., 2004). It is also feasible that Gli2 and Six1 converge on common enhancers to co-regulate target gene expression. To address this possibility, we performed ChIP-qPCR on chromatin isolated from otic vesicles at E11.5 using antibodies against Gli2 and Six1. Putative inner ear enhancers (IEEs) with Gli and Six binding sites in the vicinity of Shh responsive genes (*Frem1*, *Ece/1*, *Fam107a*, *Hey1*, *C/dn22*) demonstrated significant co-occupancy of Gli2 and Six1, with the exception of *Ece/1*, which did not show Gli2 enrichment (Fig. 6E,F). These results suggest that a subset of Shh responsive genes in the inner ear may be co-regulated by Gli2 and Six1.

### *In vivo* validation of candidate inner ear enhancers

Three candidate IEEs were selected for functional validation in a transgenic mouse reporter assay based on the presence of an ATAC-seq signal, conservation of at least one Gli binding site, and proximity to a Shh responsive gene (Fig. 7A,C,E). Candidate IEEs located in introns of *Jag1*, *P/s1* and *Brip1* demonstrated significant Gli2 enrichment, as assessed by ChIP-qPCR using chromatin isolated from otic vesicles at E11.5 (Fig. 7G). Results for Gli2 and H3K27ac occupancy at IEEs were more variable in ChIP-seq datasets from whole brain and hindbrain, respectively, possibly due to under-representation of inner ear tissue in these samples (Fig. 7A,C,E). Remarkably, each of the three IEEs generated reproducible patterns of X-gal staining in the otic vesicle of transgenic embryos, recapitulating aspects of endogenous gene expression (Fig. 7B,D,F and Table 1). ATAC-seq peaks in the vicinity of two other Shh responsive genes (*Gas2* and *Fam107a*) that did not contain Gli binding sites failed to activate reporter expression in transgenic embryos (Table 1). These results suggest that chromatin accessibility on its own may not be sufficient to accurately predict genomic regions with tissue specific enhancer activity in the inner ear. In sum, our integrated genomic approach successfully identified Shh dependent genes and enhancers in the inner ear that should assist future studies designed to address the functional impact of these factors on cochlear duct outgrowth.

**Table 1.** Results of *in vivo* mouse transgenic reporter assay for putative inner ear enhancers in the vicinity of Shh responsive genes.

| ATAC-seq peaks<br>(chromosome coordinates) | Nearest<br>gene | Gli2 ChIP-seq<br>(head) | Presence of<br>Gli motif | Transgenic<br>embryos (n) | Inner ear reporter<br>activity (n) |
| --- | --- | --- | --- | --- | --- |
| Chr2:136,932,702-136,936,730 | Jag1 | Yes | Yes | 6 | 4 |
| Chr9:95,713,880-95,717,088 | Pls1 | No | Yes | 6 | 4 |
| Chr11:85,972,172-85,973,065 | Brip1 | No | Yes | 7 | 4 |
| Chr14:9,163,756-9,164,848 | Fam107a | No | No | 9 | 0 |
| Chr7:59,140,441-59,141,435 | Gas2 | No | No | 8 | 0 |

**Figure 7.**
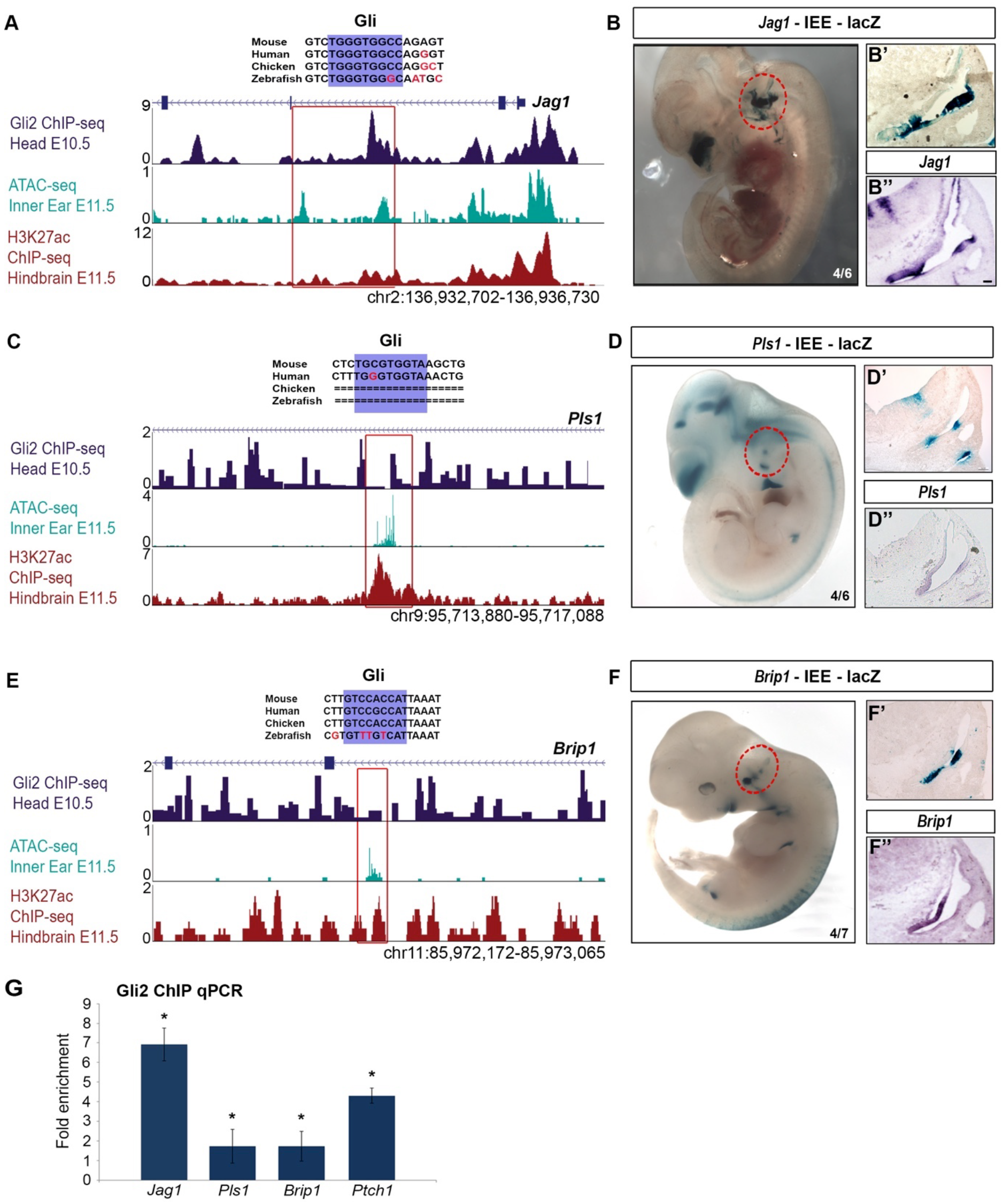
*In vivo* validation of inner ear enhancers. (A,C,E) UCSC genome browser tracks display regions of open chromatin, Gli2 binding and H3K27ac enrichment in the vicinity of Shh responsive genes (*Jag1, P/s1 and Brip1*). Boxed regions (red) represent location of inner ear enhancers. (B,D,F) X-gal staining of transgenic embryos with lacZ reporter constructs driven by IEEs. The number of embryos showing reporter activity in the otic vesicle (red circle) over the total number of transgenic embryos is depicted for each IEE. (B’, D’, F’) Transverse sections through the otic vesicle of representative transgenic embryos at E11.5 reveals a pattern of X-gal staining that is comparable to the expression of the nearby gene (B’’, D’’, F’’). (G) ChIP qPCR analysis of Gli2 binding on IEEs. A *Ptch1* enhancer is included as a positive control (*p<0.05, Student’s t-test). Error bars represent standard error of the mean.

## DISCUSSION

### Classification of Shh responsive genes during inner ear development

We exploited *Smo*^*ecko*^ and *Shh-P1* mutant embryos to identify a comprehensive set of differentially expressed genes in the inner ear that are transcriptionally activated and repressed by Shh signaling during the initial stages of cochlear duct outgrowth at E11.5. Many of these differentially expressed genes have well defined roles in cochlear development and/or auditory function but were not previously known to be dependent on Shh for their expression. Our analysis also uncovered genes with previously uncharacterized inner ear expression that represent strong candidates for follow up studies to investigate their roles in cochlear morphogenesis.

A novel outcome of this work is the observation that Shh responsive genes are partitioned into four expression domains in the ventral half of the otic vesicle. This finding reveals several new insights into the role of Shh in assigning regional and cellular identities to otic epithelial progenitors. Firstly, the ‘broad ventral’ class of Shh responsive genes distinguishes auditory (ventral) from vestibular (dorsal) regions of the otic vesicle (Fig. 2). Secondly, the downregulation of genes (e.g. *Eya1*, *Six1*, *Jag1* and *Sox2*) with essential roles in prosensory development in ‘broad ventral’, ‘medial wall’ and ‘ventral tip’ regions of the otic vesicle in *Smo*^*ecko*^ embryos highlights a previously unappreciated role for *Shh* in regulating transcription of genes that prime the medial wall of the cochlear duct for subsequent sensory development. Thirdly, loss of *Otx2* expression on the ventrolateral side of *Smo*^*ecko*^ otic vesicles likely explains the ectopic expression of prosensory markers, indicating an additional role for Shh in ensuring correct positioning of the prosensory domain. Since conditional *Otx2* mutants also show ectopic expression of prosensory markers in non-sensory regions of the cochlear duct (Vendrell et al., 2015), and that *Mycn* and *Six1* mutants display similar phenotypes, including the downregulation of *Otx2* expression (Ozaki et al., 2004; Vendrell et al., 2015), we propose that Shh/Gli, Mycn and Six1 cooperate to regulate *Otx2* expression on the ventrolateral side of the otic vesicle.

Prior studies demonstrated requirements for Shh in limiting the size of the prosensory domain and preventing precocious cell cycle exit and/or differentiation of sensory progenitors (Driver et al., 2008; Bok et al., 2013). These functions contrast with those described in our study in that they are mediated by a different source of Shh (spiral ganglia versus notochord) acting at a later stage of development (E13.5 versus E10.5). Thus, Shh signaling fulfills spatially and temporally distinct roles in regulating positive and negative aspects of sensory epithelia formation in the cochlear duct.

### Characterization of inner ear enhancers regulating Shh responsive genes

The overlap in expression of Shh responsive genes with transcription factors (*Gli1*, *Pax2*, *Gata3*, *Otx2*) in each of the four Shh responsive regions of the otic vesicle suggested a possible mode of regulation. In partial support of this claim, we observed an enrichment of Shh responsive genes in the vicinity of putative IEEs identified by sites of chromatin accessibility and Gli2 occupancy. Gli motifs were the most abundant TFBS in these putative IEEs, whereas, binding sites for other Shh dependent transcription factors (Pax2, Gata3, Otx2) were not enriched in this dataset. These results suggest that Pax2, Gata3 and Otx2 may not play a significant role in the direct regulation of Shh responsive genes at E11.5, but leaves open the possibility that they may do so at earlier stages of inner ear development.

Despite the absence of predicted TFBS from putative Shh dependent IEEs, our analysis identified other transcription factors, including Six1 and Sox2, that may cooperate with Gli2 in the direct regulation of Shh responsive genes in the otic vesicle at E11.5. Six1 is a particularly compelling candidate given the phenotypic similarities between *Six1*^−/−^ and *Smo*^*ecko*^ mutants, including cochlear agenesis and altered dorsoventral patterning of the otic vesicle, suggesting that Gli2 and Six1 may regulate common target genes (Zheng et al., 2003; Ozaki et al., 2004; Brown and Epstein, 2011). The co-recruitment of Gli2 and Six1 to a subset of putative IEEs in the vicinity of Shh responsive genes is consistent with this premise.

We also demonstrated that sites of open chromatin overlapping with conserved Gli binding sites in the vicinity of Shh responsive genes are good predictors of functional IEEs. Interestingly, IEEs with optimal (*Jag1*) and low to moderate (*Pls1, Brip1*) affinity Gli motifs each directed patterns of reporter activity in the ventral otocyst that resembled expression of the nearby gene. Thus, unlike Hh/Gli signaling in other developmental contexts, Gli binding site quality does not appear to correlate with enhancer activity in Shh responsive cells in the inner ear (Peterson et al., 2012; White et al., 2012; Ramos and Barolo 2013; Lorberbaum et al., 2016).

The three genes (*Jag1*, *P/s1* and *Brip1*) for which IEEs were identified were not previously known to be regulated by Shh signaling. Gain and loss of function studies indicate that Jag1, a Notch ligand, is required for prosensory development (Kiernan et al., 2001; Kiernan et al., 2005; Kiernan et al., 2006; Brooker et al., 2006). The initiation of *Jag1* expression in the otic placode is regulated by Wnt signaling (Jayasena et al., 2008), whereas our data suggests that maintenance of *Jag1* in prosensory progenitors is dependent on Shh. Pls1 is an actin bundling protein that maintains the length and width of stereocilia in inner hair cells and is required for optimal hearing in adult mice (Taylor et al., 2015). Since Pls1 is dispensable for the initial formation of stereocilia, it remains to be determined what role its Shh regulated expression might play during otic development. Similarly, the inner ear function of Brip1, a member of the RecQ DEAH helicase family that interacts with Brca1 in DNA damage repair and tumor suppression, is currently unknown (Ouhtit et al., 2016).

In summary, our integrated genomic approach greatly expands the list of genes and regulatory sequences that depend on Shh signaling in the inner ear. These datasets should benefit future studies addressing the function and regulation of key genes acting downstream of Shh that promote cochlear duct morphogenesis and establish its distinct cellular composition.

## MATERIALS AND METHODS

### Mouse Lines

All mouse experiments were performed in accordance with the ethical guidelines of the National Institutes of Health and with the approval of the Institutional Animal Care and Use Committee of the University of Pennsylvania. The production of *Smo*^*ecko*^ (*Foxg1cre; Smo*^*loxp/−*^) and control (*Foxg1cre; Smo*^*loxp/+*^) embryos was described previously (Brown and Epstein, 2011). The *Shh-P1* mouse line was described previously (Riccomagno et al., 2002).

### RNA-seq analysis

Otic vesicles from control, *Smo*^*ecko*^ and *Shh-P1* embryos (n=4 pairs of biological replicates for each genotype) were isolated at E11.5, exposed to collagenase P (1mg/ml) at 37°C for 20 min to remove surrounding mesenchyme, and submerged in RNA*later*™ Stabilization Solution (Thermo Fisher Scientific, Cat#AM7022). RNA was extracted using the RNeasy Micro Kit (Qiagen, Cat#74004). Total RNA (200ng) was used for poly A selected RNA-seq library preparation using the NEBNext® Ultra™ Directional RNA Library Prep Kit for Illumina® (mRNA) (New England Biolabs Cat#E7530S). Biological replicates were individually barcoded, pooled, and sequenced to generate 100bp single-end reads on one lane of a HiSeq4000 instrument at the Next Generation Sequencing Core (Perelman School of Medicine, University of Pennsylvania). RNA-seq reads were aligned to the mm9 mouse genome build (http://genome.ucsc.edu/) using RUM (Grant et al. 2011). Differential gene expression analysis between *Smo*^*ecko*^, *Shh-P1* and control samples was performed using edgeR (Robinson et al., 2010). Heatmaps for RNA-seq data were generated using PIVOT (Zhu et al., 2018). RNA-seq data were deposited in NCBI GEO under accession number GSE131165.

### ATAC-seq analysis

Four independent ATAC-seq libraries were generated from wild type otic vesicles (n=10 per library) isolated at E11.5 using 50,000 cells per replicate as described (Buenrostro et al., 2015). Tagmentation was performed using the Nextera® DNA Library Preparation Kit (lllumina® 15028211). Multiplexed 50 bp paired-end sequence reads were generated on a single lane of an lllumina HiSeq2000 instrument. ATAC-seq reads were mapped to the mm9 mouse genome build (http://genome.ucsc.edu/) using Bowtie with default parameters (Langmead et al., 2009). Regions of open chromatin were identified by MACS2 using default parameters (Zhang et al., 2008). Only high confidence peaks that were present in at least two libraries were reported. ATAC-seq data were deposited in NCBl GEO under accession number GSE131165.

### Gli2 ChIP-seq

Mouse embryonic tissues were harvested from timed matings between Swiss Webster (Taconic) mice where day of detection of vaginal plug was considered embryonic day E0.5. ChIP was performed on pooled tissue obtained from 40 E10.5 heads isolated below the otic vesicle and processed for ChIP as described (Peterson et al., 2012). Briefly, embryonic tissue was fixed for 30 minutes in 1% formaldehyde/PBS at room temperature followed by quenching with 125 mM glycine. ChIP was performed on the entire lysate using magnetic Dynabeads^TM^ Protein G (Thermo Fisher Scientific) bound with goat anti-Gli2 antibody (R&D Cat# AF3635). A ChIP DNA library was prepared for lllumina Sequencing according to manufacturer recommendations and 50 bp single-end reads were obtained from a Hi-Seq2000 instrument. The resulting reads were mapped to mouse genome assembly mm9 (http://genome.ucsc.edu/) using Bowtie (Langmead et al., 2009). Gli2 ChIP-seq data were deposited in NCBl GEO under accession number GSE131165.

### Intersection of ATAC-seq and Gli2 Ch/P-seq data

Enriched peaks from the ATAC-seq and Gli2 ChIP-seq were intersected using Bedtools intersect interval function (Galaxy Version 2.27.1+galaxy1) with the parameter, *-wa, overlap on either strand* and returning full length ATAC-seq peaks that overlap with Gli2 ChIP-seq peaks (Quinlan and Hall, 2010).

### ATAC-seq enrichment at inner ear promoters and VIS1A enhancers

ATAC-seq promoter peaks were first identified by intersecting RefSeq (mm9) genes extracted from UCSC Genome Browser (Table Browser; parameter: create one Bed record per upstream by 500 bp) using Bedtools intersect interval function (Galaxy Version 2.27.1+galaxy1) with the parameter, *-wa, overlap on either strand* and returning RefSeq peaks. The gene name was then used to compare against E11.5 control RNA-seq genes that are expressed (RPKM≥1.0) or not expressed (RPKM<1.0) in the otic vesicle. The enrichment of ATAC-seq peaks against VISTA enhancers (https://enhancer.lbl.gov/) was performed by converting enhancer coordinates to peak files and then intersected with ATAC-seq non-coding peaks using Bedtools (parameters similar to above and returning ATAC-seq peaks).

### ATAC-seq conservation analysis

We measured placental mammal-derived (PS) and vertebrate-derived (VS) PhyloP scores within each ATAC-seq peak, and estimated the ratio of these values for each tissue. For each tissue, we first identified the mode of the enhancer peak width distribution, so that each called peak could be elongated or trimmed to a standard peak size. Then, extending peaks shorter than this mode, or contracting peaks longer than the mode, we estimated PS and VS for each base of each peak. For each tissue, the mean PS and VS were higher at the center of the peak compared to the edges. Since the PS and VS have different ranges of values, we used their ratio for comparison between tissues instead of using their absolute values. For each base position, we then obtained the ratio of the mean PS score to the mean VS score across all peaks in that tissue. As a control, we also took the set of all exons in the mouse genome, and defining a peak of length 201, obtained PS and VS for each base of each exon.

### Functional annotation analysis of ATAC-seq data

Functional annotation analysis of ATAC-seq data was performed using GREAT version 3.0.0 (McLean et al., 2010), linking peaks to the nearest transcription start site (TSS) ± 100 kb. Functional terms were selected based on reported significance score and relevance to the biological system.

### Gene set enrichment analysis (GSEA)

GSEA was performed using the GSEA software (MSigDB 6.1 and 6.2) as described (Mootha et al., 2003; Subramanian et al., 2005). Ranked file (rnk) for Shh activated genes was prepared from the RNA-seq differential expression analyses based on the log_2_ fold change. The. grp files were prepared from expressed genes (i.e. RNA-seq, RPKM ≥ 1.0) that are found within the intersected peaks: i) ATAC-seq and Gli2 ChIP-seq ± 500 kb (Fig. 6C) and ii) ATAC-seq, Gli2 ChIP-seq and H3K27ac ChIP-seq peaks ± 500 kb (Fig. S4).

### Ch/P-qPCR

Otic vesicles were dissected in DMEM (with 10% fetal bovine serum) from approximately 25-30 E11.5 embryos per replicate pool (n=3 replicates), homogenized into small pieces, and crosslinked with 1% paraformaldehyde for 15 min at room temperature with shaking. ChIP was performed essentially as described on three biological replicates (Zhao et al., 2012) using 6 µg of anti-Gli2 (R&D Cat# AF3635), anti-Six1 (Cell Signaling Technology, Cat#12891) or anti-immunoglobulin G (lgG) (Cell Signaling Technology) antibodies. QPCR was conducted as described (Zhao et al., 2012) using primer sequences listed in Table S5. Positive control primers in Fig. 7G amplify a DNA fragment from a *Ptch1* enhancer bound by Gli2.

### Transgenic mouse reporter assay

Candidate inner ear enhancers were cloned into a vector containing the *Hsp68* promoter, /*acZ* gene and SV40 poly(A) cassette. Transient transgenic embryos were generated by pronuclear injection into fertilized mouse eggs derived from the (BL6xSJL) F1 mouse strain (Jackson Laboratories) at the Transgenic and Chimeric Mouse Facility (Perelman School of Medicine, University of Pennsylvania). For X-gal staining, embryos were harvested at E11.5, fixed in 0.2% glutaraldehyde/1% formaldehyde at 4°C for 30 minutes, and stained in a solution containing 1 mg/ml X-gal at 37°C for two hours to overnight.

### Statistical analysis

Relevant information for each experiment including n-values, statistical tests and reported p-values are found in the legend corresponding to each figure. In all cases ps0.05 is considered statistically significant and error bars represent standard error of the mean.

### Data availability

RNA-seq, ATAC-seq and ChIP-seq datasets have been deposited in NCBI under GEO accession number GSE131165.

## Acknowledgements

We thank Dr. Jean Richa and his staff at the Transgenic and Chimeric Mouse Facility (Perelman School of Medicine, University of Pennsylvania) for their assistance in transgenic mouse production, as well as Dr. Jonathan Schug and his team at PSOM NGSC for sequencing services. We also thank members of the Epstein lab for critical comments on this study.

## Competing Interests

No competing interests declared.

## Author contributions

V.M., A.M.R., S.M.R., Y.Y., A.S.B., and K.A.P performed the experiments. Y-T.Z., J.M., K-J.W., S.R., C.D.B. and V.M. performed data analysis, V.M., Y.Y. and D.J.E. conceived the project. V.M. and D.J.E. wrote the manuscript.

## Funding

This work was supported by a grant from the National Institutes of Health [R01 DC006254] to D.J.E.

